# Optical photothermal infrared imaging using metabolic probes in biological systems

**DOI:** 10.1101/2024.09.19.613881

**Authors:** Sydney O. Shuster, Anna E. Curtis, Caitlin M. Davis

## Abstract

Infrared spectroscopy is a powerful tool for identifying biomolecules. In biological systems, infrared spectra provide information on structure, reaction mechanisms, and conformational change of biomolecules. However, the promise of applying infrared imaging to biological systems has been hampered by low spatial resolution and the overwhelming water background arising from the aqueous nature of in cell and *in vivo* work. Recently, optical photothermal infrared microscopy (OPTIR) has overcome these barriers and achieved both spatially and spectrally resolved images of live cells and organisms. Here, we determine the most effective modes of collection on a commercial OPTIR microscope for work in biological samples. We examine three cell lines (Huh-7, differentiated 3T3-L1, and U2OS) and three organisms (*E. coli*, tardigrades, and zebrafish). Our results suggest that the information provided by multifrequency imaging is comparable to hyperspectral imaging while reducing imaging times twenty-fold. We also explore the utility of IR active probes for OPTIR using global and site-specific noncanonical azide containing amino acid probes of proteins. We find that photoreactive IR probes are not compatible with OPTIR. We demonstrate live imaging of cells in buffers with water. ^13^C glucose metabolism monitored in live fat cells and *E. coli* highlights that the same probe may be used in different pathways. Further we demonstrate that some drugs (e.g. neratinib) have IR active moieties that can be imaged by OPTIR. Our findings illustrate the versatility of OPTIR, and together, provide a direction for future dynamic imaging of living cells and organisms.

## INTRODUCTION

Vibrational microspectroscopy has shown significant promise as a technique for understanding biophysical and biological processes. Infrared and Raman based techniques can provide detailed insight for complex sample identification without the need for bulky, disruptive labels.^1–4^ Vibrational techniques can also be used to assign protein secondary structure,^5^ observe changes in folding,^6^ and follow metabolic processes using isotope labels and other vibrational tags.^3^ Yet, use of vibrational imaging has been limited in cell biology and biophysics. Raman imaging suffers from weak cross-sections necessitating high laser powers. Stimulated Raman scattering has been invaluable due to its increased signal intensity, but collecting full spectrum is nontrivial, limiting its use.^4,7^ Infrared imaging, on the other hand, has not been used significantly outside ‘stainless histology’ and dry, fixed cell work because of the strong background absorption from water and poor diffraction limits, though many infrared metabolic probes have been developed^3,8,9^

Optical photothermal infrared microscopy (OPTIR), also known as mid-infrared photothermal imaging (MIP), is a new and promising technique in the biological and chemical sciences.^10–13^ OPTIR is a pump-probe technique in which a sample is pumped with a pulsed, tunable infrared laser.^10,11^ Upon absorption of the infrared light, the sample heats according to the photothermal effect. This heating causes thermal expansion and a change in the refractive index of the sample which can be probed by a visible laser. By detecting the visible laser, changes in its intensity that match the pulse rate of the infrared laser can be extracted and used to generate an infrared spectrum that corresponds well with the FTIR spectrum. This technique is analogous to AFM-IR with an optical probe instead of a physical probe.

OPTIR brings significant benefits to biophysics and biology. As a vibrational method, OPTIR allows for detailed spectral information on samples including label free identification of biopolymers and assignments of protein structures.^14^ Low laser powers are non-perturbative to the system.^15^ According to Onsanger’s fluctuation dissipation theorem^16^, the kinetics following a small perturbation away from equilibrium are identical to the natural kinetics observed by spontaneous fluctuations. Therefore, small <3 K perturbations introduced by OPTIR lasers mimic native fluctuations.^15^ Furthermore, OPTIR has a spatial resolution of under 500 nm and is less impacted by water absorption as water’s high heat capacity reduces its photothermal signal. Because of this, OPTIR can be used to collect vibrational images in living cells and tissues in water. Past work has demonstrated that OPTIR can be used to collect both single frequency images and full spectra in live cells.^12,17,18^ Additionally, hyperspectral images have been collected in fixed and dried tissues.^13,17^ These techniques have been used to track lipid^15,17,18^ and carbohydrate^19^ metabolism in cells and identify protein aggregation in tissues.^20^

Vibrational probes used with OPTIR include isotope labels, such as ^13^C and ^2^H, and small IR active moieties, such as azides and nitriles. Carbon-deuterium, azide, and nitrile moieties absorb at ∼2100 cm^-1^ which falls into the “cellular silent region” where other cellular biomolecules do not absorb allowing for easy discernment of the labeled substrate from cellular background. This region has been used with significant success in spontaneous Raman^21^ and SRS^22^ with alkyne moieties.

OPTIR has the potential to be most informative in living cells, tissues, and organisms where cellular processes could be measured in a tag-free manner or using small IR probes. Additionally, OPTIR intended for biological systems is now commercially available via the mIRage-LS (life science) from Photothermal, Inc. Although OPTIR shows significant promise, it has been limited by slower acquisition times (in the case of hyperspectral images) and reduced spectral information that may not well correlate to full spectra (in the case of single frequency images). Here, we evaluate the commercial instrument and examine corrected single frequency imaging and multispectral ratio imaging as compared to hyperspectral imaging of fixed cells to assess the accuracy of multispectral imaging. We conclude that when properly corrected for and used for ratio imaging, multispectral images can provide similar information to hyperspectral images with 1/20^th^ of the collection time. Importantly, we show that due to the use of visible lasers, not all IR active probes are OPTIR compatible. Nevertheless, we highlight the versatility of the technique, performed on a commercial instrument, with various IR probes of live cell lines and model organisms imaged in water. This lays the groundwork for future OPTIR studies of metabolic processes in live cells and organisms.

## METHODS

### Materials

Unless otherwise specified, all reagents were sourced from Sigma-Aldrich.

### Oleic Acid Conjugation and Feeding Protocol

Oleic acid conjugation and feeding was performed as previously described.^18^ Briefly, ^2^H oleic acid (oleic acid-d_33_, DLM-1891-PK, Cambridge Isotope Laboratories, Tewksbury, MA) was bound to fatty acid-free bovine serum albumin (BSA) using a protocol from Shi et al.^3^ The stock oleic acid concentration is approximately 3 mM. The ratio of oleic acid to BSA is 2:1, in line with physiological ratios.^23,24^ Oleic acid is then incubated with cell culture at a concentration of 60 µM.

### Mammalian cell culture

Huh-7 cells (gift from Lars Plate, Vanderbilt University) and U2OS cells (ATCC, Manassas, VA) were prepared as previously described.^17,18^ Briefly, cells were cultured in 75 cm^3^ sterile vented cap tissue culture flasks (Corning, Corning, NY). At 80 % confluence, cells were trypsinized and replated on CaF_2_ coverslips (20 × 20 × 0.35 mm, Crystran, Poole, U.K.) in 35 mm diameter sterile Petri dishes (Corning) for live samples and CaF_2_ coverslips (10 × 0.35 mm) in a 24 well cell culture plate (Corning) for fixed samples. Cells were allowed to adhere for 24 hours prior to experimentation.

3T3-L1 cells (ATCC, Manassas, VA) were prepared as previously described.^17^ Briefly, low passage number (<10), predifferentiated cells were grown to confluence in Dulbecco’s modified Eagle’s medium containing 4.5 g/L glucose and Lglutamine (DMEM, Corning) supplemented with 10% calf bovine serum (CBS, ATCC) and 1% penicillin/streptomycin (Thermo-Fischer) under standard conditions. Two to three days post confluence the cells were differentiated, media was changed to DMEM containing 10% fetal bovine serum (FBS, Corning), 1% penicillin/streptomycin, 20 μg/mL insulin, 250 nM dexamethasone, and 500 μM isobutylmethylxanthine. Two to three days post differentiation, media was changed to DMEM containing 10% fetal bovine serum (FBS, Corning), 1% penicillin/streptomycin and 20 μg/mL insulin. Following another two to three days, cells were trypsinized and replated on CaF_2_ coverslips (20 × 20 × 0.35 mm; Crystran.) in 35 mm dishes (Corning). Media was exchanged every two to three days until lipid droplets formed and stabilized (five to seven days).

### ^13^C and ^2^H labeling mammalian cells

Medium was replaced with glucose-free DMEM supplemented with 1% FBS, 1% penicillin/streptomycin, ^13^C glucose (D-glucose-^13^C_6_, CLM-1396-PK, Cambridge Isotope Laboratories) to 4.5 mg/mL, and either 2% BSA or 2% ^2^H oleic acid conjugated with BSA (final concentration 60 µM oleic acid and 30 µM BSA).

### Neratinib treatment of U2-OS cells

Cells were treated with 5 uM neratinib (Cayman Chemical, Ann Arbor, MI) for 7 hours and then imaged live.

### ^13^C labeling *E. coli*

BL21(DE3) strain *E. coli* were transformed with pET28a-sfGFP, gift from Ryan Mehl (Addgene plasmid #85492). Several colonies were selected and grown in 5 mL of Luria broth (LB) at 37 °C with 300 rpm shaking overnight. 200 μL of starter was transferred to 5 mL of M9 media supplemented with ^13^C labeled glucose or LB (control) and grown until cloudy, then 1 mL was moved to 20 mL of M9 supplemented with ^13^C labeled glucose or LB (control) and grown until an OD at 600 nm of 0.6-0.8. The temperature was then reduced to 20 °C and the *E. coli* grown for 16 hours.

### Mammalian cell fixing

For fixed cell samples, mammalian cells were fixed at the desired time points with 4% paraformaldehyde (PFA) in PBS (Corning) for 20 minutes at room temperature. The fixed cells were washed three times with PBS followed by three times with Milli-Q purified water.

### Live cell chamber construction

Live samples cultured on CaF_2_ coverslips were washed once with PBS and mounted in PBS or Fluorobrite DMEM (Gibco) on a glass microscopy slide (VWR). A double-sided tape spacer (Nitto, San Diego, CA) with a 5 μm thickness was used to keep the cells hydrated and minimize compression of the cell.

### Azide incorporation in E. Coli

The azidohomoalanine (AHA) incorporation was adapted from homopropargylglycine incorporation protocols.^1^ B834 (DE3) methionine auxotroph *E. coli* (Novagen) were transformed with pUC19. A single colony was used to seed 5 mL of LB and grown overnight at 37 °C with shaking at 300 rpm. In the morning, 200 μL of the LB starter was used to seed 5 mL of M9 media supplemented with ampicillin (100 µg/mL) and methionine (40 µg/mL) and grown with shaking at 300 rpm at 37 °C until cloudy (∼4 hours). One mL of this was used to seed 20 mL of M9 media supplemented with ampicillin (100 µg/mL) and methionine (40 µg/mL) and grown with shaking at 300 rpm at 37 °C. At OD600 ∼0.6-0.8, the sample was centrifuged at 3900 rpm for 20 minutes. The resulting pellet was resuspended in 20 mL fresh M9 and starved at 37 °C with 300 rpm shaking for 30 minutes. Ampicillin (100 µg/mL) and AHA (40 µg/mL) were supplemented and the temperature reduced to 20 °C for an overnight expression.

### 4-azidomethyl-L-phenylalanine (AzmF) and 4-azido-Lphenylalanine (azF) incorporation

AzmF and AzF incorporation was adapted from Bazewicz et al.^25^ B.95ΔAΔfabR *E. coli*^26^ (Yale *E. coli* Genetic Stock Center, New Haven, CT) were made chemically competent^27^ and co-transformed with pET28a-sfGFP TAG150 and pDule-pCNF, gifts from Ryan Mehl (Addgene plasmid #85493 and #85494).^28^ A single colony was selected and used to inoculate 5 mL of LB. The starter was grown overnight at 37 °C and 300 rpm shaking. 1 mL of the starter was used to inoculate 20 mL of LB with kanamycin (50 µg/mL) and tetracycline (10 µg/mL) and grown at 37 °C until OD600 ∼0.5-0.6 when AzmF or AzF (Chem Impex, Wood Dale, IL) was added to 1 mM and IPTG to 1 mM. The sample was kept at 20 °C with shaking at 300 rpm for 16 hours.

### Single and bulk *E. coli* preparation

For bulk analysis a 1 mL aliquot of E. coli was harvested at 3900 rpm and the pellet collected and dried on a glass slide. For fixed and live single cell preparation a 1 mL aliquot of *E. coli* were washed three times in PBS. For live samples, *E. coli* was resuspended in 1 mL PBS. 30 μL was added to a 5 uM spacer on a glass slide. The chamber was closed with a 1% polyethyleneimine (PEI) coated CaF2 coverslip. For fixed samples, *E. coli* was fixed with 4% PFA for 20 minutes and then washed three times with PBS and two times with Milli-Q H2O. The final pellet was resuspended in 100 μL of H2O and 10 μL spotted onto 1% PEI coated CaF2 coverslips for bulk data collection. For single cells, dilutions were made with H2O (1 mL H_2_O was generally sufficient) and then spotted onto a 1% PEI coated CaF_2_ coverslip. Fixed samples were allowed to dry, protected from light, for 20 minutes and then imaged directly.

### OPTIR data collection

All imaging was performed as previously described^18^ on a mIRage-LS IR microscope (Photothermal Spectroscopy Corporation, Santa Barbara, CA) integrated with a four-module-pulsed quantum cascade laser (QCL) system (Daylight Solutions, San Diego, CA) with a tunable range from 932 cm^-1^ to 2348 cm^-1^ unless otherwise specified. Briefly, brightfield optical images were collected using a low magnification 10X refractive objective with a working distance of 15 mm. Spectra and infrared images were collected in copropagating mode unless otherwise specified using a 40X Cassegrain objective with a working distance of 8 mm. Fixed cell spectra and images were collected in standard (epi) mode and live cell spectra and images were collected in transmission mode. Data was collected with an IR laser power of 20% and a probe power of 11%. Laser power was selected to maximize signal while minimizing sample damage and heating. For live cells under our collection conditions, we have AC values between 5 and 50 mV with a DC voltage of ∼3.5 V and a detector gain of 5x corresponding to temperature changes between 0.3 and 3 K. All spectra and images were collected using PTIR Studio 4.5 (Photothermal Spectroscopy Corporation). Image acquisition time for multispectral images was 2-5 minutes per frequency for a total acquisition time of 10-25 minutes per cell. For hyperspectral images, a step size of 500 nm, the maximum resolution of the instrument, and one acquisition was used.

### Fluorescence imaging

Fluorescent images were collected using the mIRage-LS, which is equipped with a Prime BSI Express sCMOS (Teledyne, Thousand Oaks, CA). Fluorescent proteins were excited using a Sola Light Engine (Lumencor, Beaverton, OR) and GFP filter cube (TLV-U-FF-GFP, Thorlabs, Newton, NJ). Images were collected with the 50X objective, 10% lamp power and 100 μs acquisition time.

### Tardigrade imaging

*H. exemplaris* tardigrades (Carolina Biological Supply, Burling, NC) were cultured in spring water (Poland Spring, Poland, ME) according to published tardigrade culturing protocol.^29^ Single tardigrades were collected using a 200 uL pipette tip and deposited onto a glass microscopy slide (VWR) in 30 uL of spring water with double-sided tape spacers (Nitto) with a 5 μm thickness. Tardigrades were covered with a 350 µm thick CaF_2_ coverslip and imaged immediately.

### Data analysis

Images were processed in Fiji (NIH, Bethesda, MD).^30^ Spectra were analyzed in IGOR Pro 9 (Wavemetrics, Portland, OR). Live and fixed cell ratio images for both multispectral^18^ and hyperspectral^17^ images were generated in Python 3.10 in Colab (Google, Mountain View, CA)^31^ as previously described.^17,18^ To account for overlapping signals of other biomolecules, the ratio images were corrected using the correction protocol adapted from Shi et al.^3^ This protocol removes contribution from the overlap of the broad amide I and/or water bands (1655 cm^-1^) to the lipid bands by subtracting the average percentage of this band (correction value) at the lipid position (^13^C=O, 1703 cm^-1^, or ^12^C=O, 1747 cm^-1^) from the observed intensity at each frequency; data collected in control samples without overlapping bands (e.g. cells not fed ^13^C or water alone) across all time points are averaged to create a correction value (b, c, d and f below) with which to subtract off overlapping amide-I and/or water from lipid intensities.^18^ By adjusting the frequencies, this procedure can be modified to correct for overlapping spectral features in any region of interest. This procedure produces a ratio that more accurately reflects the relative populations of the spectral features of interest and is crucial to normalize ratios in cells with highly fluctuating backgrounds (e.g. water in live cells).

Hyperspectral ratio images were corrected with Equation

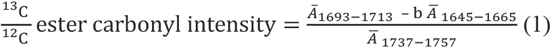

where *Ā* is the average signal intensity over the indicated frequencies. *Ā* was calculated from full spectra of fixed, control adipocytes. The variable *b* is the correction value for the amideI intensity, with *b* being 0.31 for adipocytes. Fixed cell ratios (eq. 1) were only performed on regions of the cell that had lipid, defined as a ^12^C lipid ester carbonyl intensity of 0.025 or greater after vector normalization.

Fixed cell ^13^C=O/^12^C=O multispectral ratio images were corrected for the amide-I intensity using Equation 2:

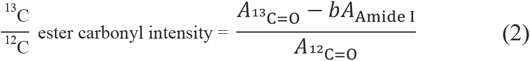

where *A* is the signal intensity of the frequency of the indicated biomolecule. The ^12^C=O stretch is at 1747 cm^-1^ for adipocytes. The ^13^C=O stretch is at 1703 cm^-1^. The variable *b* is the correction value for the amide-I intensity, with *b* being 0.31 for adipocytes. Fixed cell ratios were only performed on regions with sufficient ^12^C=O signal intensity, defined as regions with 5% of the maximum value of ^12^C=O signal for adipocytes.

Live cell ^2^H/^12^C ratio images were corrected for the water band using equation 3.

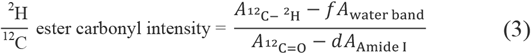

The ^12^C -^2^H band is measure from a shoulder at 2212 cm^-1^ for adipocytes. The water band is measured from 2050 cm^-1^ for both cell lines as this does not overlap with other ^12^C -^2^H stretches. The variable *f* is the correction value for the water band and is 0.85 for adipocytes and *d* is the correction value for 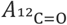 and is 0.28 for adipocytes. Live cell ratios were also only performed on regions with sufficient ^12^C lipid ester carbonyl signal intensity, defined as regions with 5% of the maximum value of ^12^C lipid ester carbonyl signal for adipocytes.

Live cell ^13^C=O/^12^C=O ratio images were corrected for the amide I and water band intensity using equation 4:

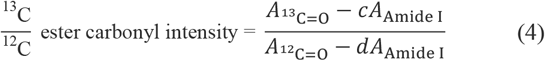

where *c* is the correction value for 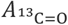 and is 0.66 for adipocytes and *d* is the correction value in equation 3. Live cell ^2^H/^12^C ratio images were corrected for the water band using equation 4.

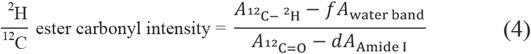

The ^12^C -^2^H is measure from a shoulder at 2212 cm^-1^ for adipocytes. The water band is measured from 2050 cm^-1^ for both cell lines as this does not overlap with other ^12^C -^2^H stretches. The variable *f* is the correction value for the water band and is 0.85 for adipocytes and *d* is the correction value discussed above. Live cell ratios were also only performed on regions with sufficient ^12^C lipid ester carbonyl signal intensity, defined as regions with 5% of the maximum value of ^12^C lipid ester carbonyl signal for adipocytes.

Other ratio images were thresholded as noted and are simple ratios of the relevant frequencies, without correction.

## RESULTS AND DISCUSSION

### Data collection methods and suitability

Our OPTIR instrument, a commercially available mIRage-LS from Photothermal Spectroscopy Corporation, can collect data in two primary formats: full spectra and single frequency imaging. The spectral coverage of the instrument can be customized by the selected IR laser; our instrument covers a range of 932 cm^-1^ to 2348 cm^-1^ with a gap between 1816 cm^-1^ and 1992 cm^-1^. Full spectra (Fig. 1A) can be collected in an ad-hoc fashion or using an array. Using an array enables the construction of hyperspectral images wherein each pixel of the image contains a full IR spectrum (Fig. 1B). Alternatively, an array can be used to collect images at a single frequency or a few frequencies of interest (multispectral images) (Fig. 1C). Multispectral images are generated by either collecting single frequency images in sequence and stacking them together or, more recently introduced in the commercial system, by an interleaving method where a row of pixels is collected at one frequency and then again at each additional frequency before imaging the next row of pixels.

**Figure 1.**
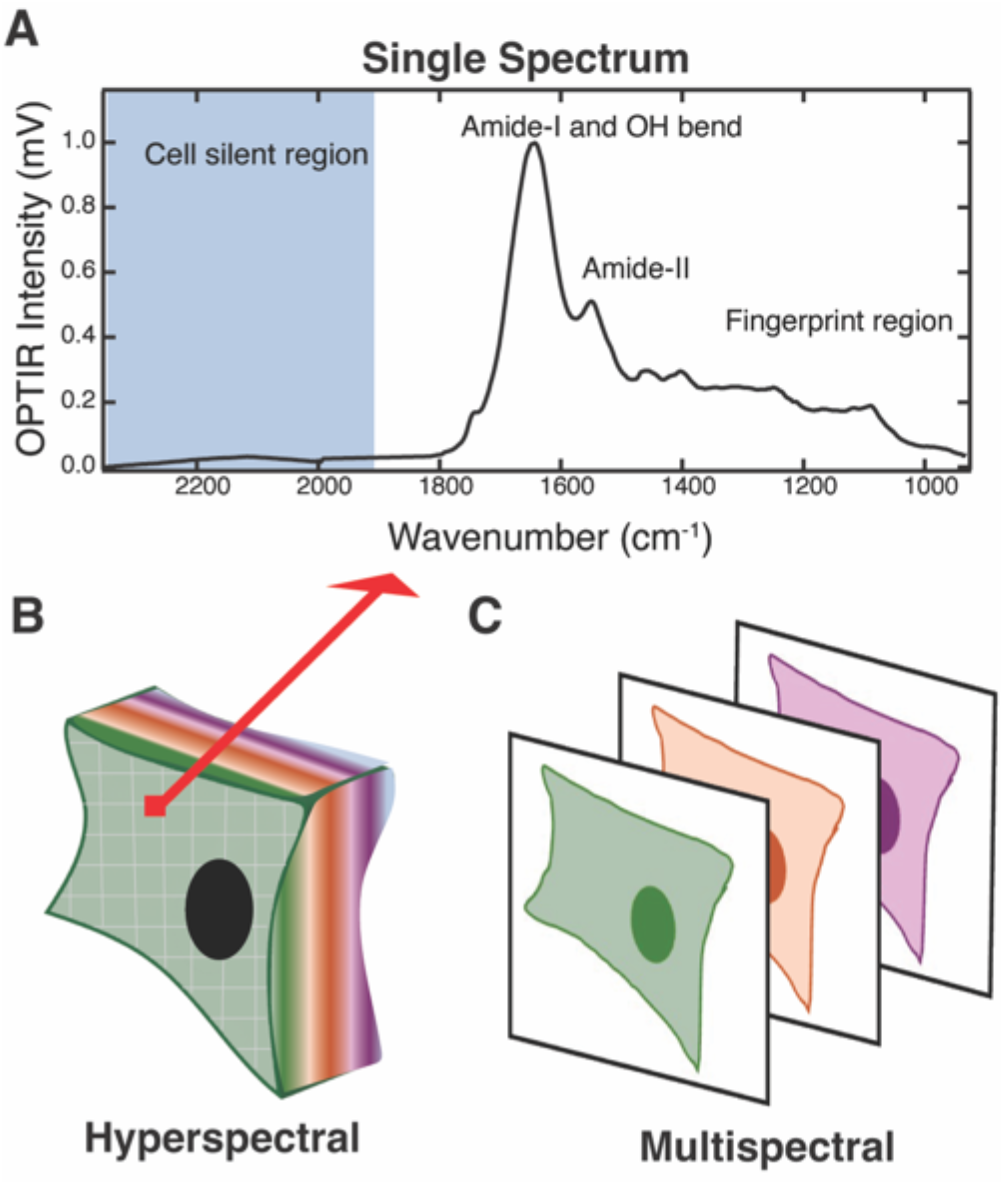
Hyperspectral vs. multispectral imaging. (A) Representative spectrum of a hydrated biological sample. The cell silent region is shown in blue. (B) Hyperspectral OPTIR map of a cell, where a full IR spectrum is collected in each 500 × 500 nm pixel. (C) Multispectral imaging where an image is collected at several discrete IR frequencies of interest.

OPTIR is compatible with a variety of organisms, cells, and tissue types.^12^ Notable IR bands in a generic biological sample (Fig. 1A) include the water bend (1640 cm^-1^, broad), the water bend-libation combination band (2150 cm^-1^, broad, weak), the amide-I (1615-1690 cm^-1^) and amide-II (∼1550 cm^-1^) of proteins, the CH deformation of proteins and lipids (1300-1500 cm^1^), nucleic acid and lipid phosphate stretches (∼1080 cm^-1^), and others.^5,32,33^ Using these bands, different cell types can be distinguished.^34–36^ We compared average fixed cell spectra collected in epi copropagating mode of full hyperspectral maps of three commonly used model cell lines: osteosarcoma line U2OS, hepatoma line Huh-7, and adipocyte model line differentiated 3T3-L1, and made full assignments (Fig. S1). The average spectra were similar with near identical amide-I and amide-II bandwidths and ratios. Although previous work^34^ has shown differences in amide-I to amide-II ratios and bandwidths between cell lines, that work distinguished cancerous and benign cells from tissue while our samples were all cultured cancer derived cell lines. Within the cancer derived lines (Fig. S1) the ratio of lipid bands as compared to the protein bands was dramatically different. Differentiated 3T3-L1 cells, which are an adipocyte-like cell lines and therefore lipid rich, had the highest lipid to protein ratio. U2OS cells had the lowest. There were also notable differences in the CH deformation region, pointing at differences in protein, lipid, and carbohydrate composition.

To determine the optimal mode for imaging biological samples, the signal-to-noise (S/N) ratio of the different OPTIR imaging modalities was evaluated. The OPTIR instrument’s laser paths can be set up in two modes. In copropagating mode (coprop, Fig. 2A), the infrared and visible lasers are collinear. The visible laser can be collected for analysis either in epi or transmission mode. In counterpropagating mode (counterprop, Fig. 2B), the visible and infrared lasers are split with the visible laser entering from the top objective and the infrared from the bottom objective. This can again be collected in epi or transmission mode. The noise of each modality was determined by collecting three reference spectra of a calcium fluoride coverslip, which has no mid-IR absorbance, for each modality (Fig. S2).

**Figure 2.**
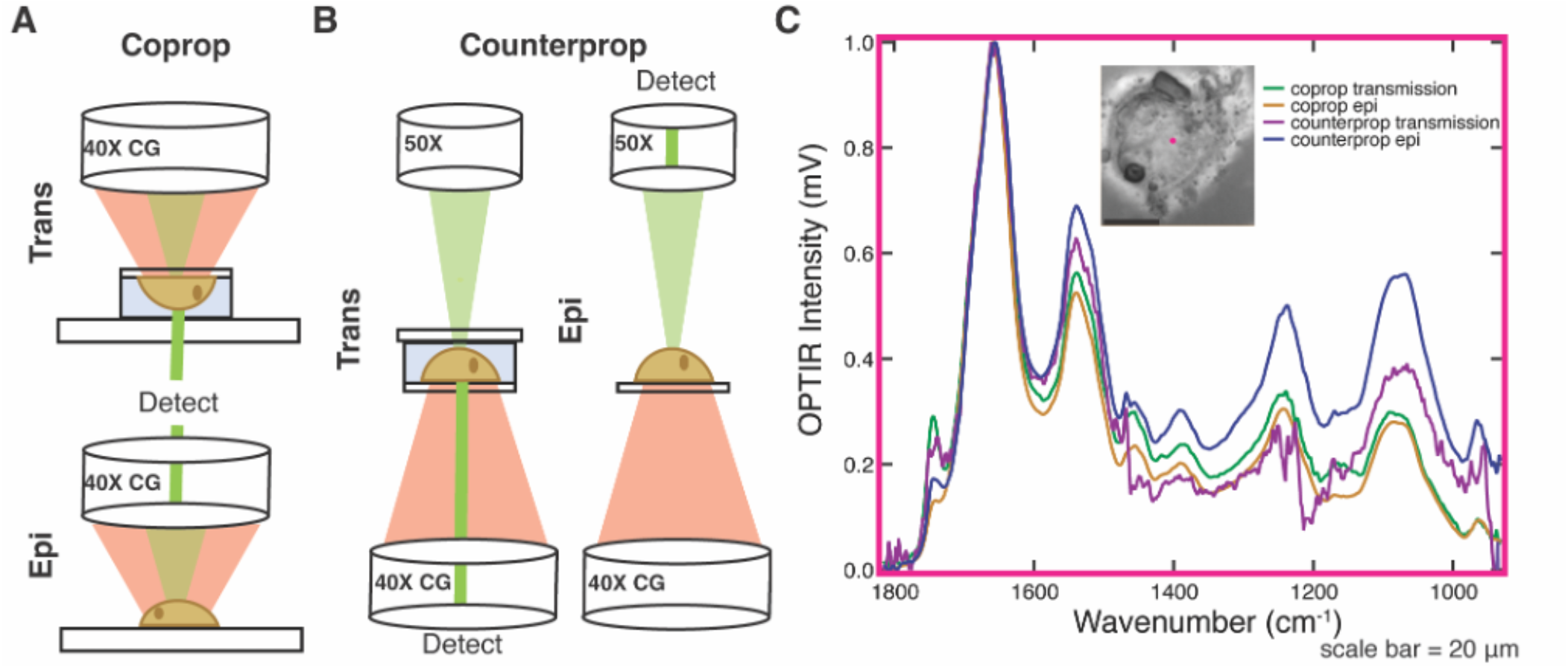
Signal-to-noise (S/N) ratio of different modes of data collection. The visible laser of the OPTIR can be collected in both and epi and trans geometry for both (A) copropagation and (B) counterpropagation geometry of the infrared laser. Trans geometries show a live cell in an aqueous chamber but can also be collected on a fixed sample. (C) Example spectra from a fixed and dried Huh-7 cell collected at the same spot (x, y, and z held constant, pink) in coprop transmission (green) and epi (yellow) and counterprop transmission (purple) and epi (blue).

Each spectrum is the average of three acquisitions with an IR power of 20%, probe power of 11%, and detector gain of 1x. We report average root mean square noise across covered frequencies: 0.02 mV for coprop transmission, 0.01 mV for coprop epi, 0.31 mV for counterprop transmission and 0.04 mV for counterprop epi. Smaller signals can be detected by additional signal averaging. Using a fixed and dried cell grown on CaF_2_ (Fig. 2C) as a representative sample gives S/N ratios at 1658 cm^-1^ of ∼514 for coprop trans, ∼2760 for coprop epi, ∼33 for counterprop trans, and ∼1150 for counterprop epi. In all modalities, the S/N ratio of coprop is greater than counterprop, and this difference can be visualized in spectra of dried, fixed cells across the spectral range (Fig. 2C). Because of the higher S/N ratio, we commonly collect data in coprop mode. However, counterprop is not without advantages as it has a slightly improved spatial resolution. This is due to the improved numerical aperture of the 50x refractive objective (NA 0.8) used for the visible light in counterprop (Fig. 2B) as compared to the 40X Cassegrain objective (NA 0.78) used for the infrared/visible light (Fig. 2A). It also resolves minor issues caused by changing between objectives when comparing brightfield or fluorescence images to OPTIR images.

While epi is generally used for dried, fixed cell work, transmission is compatible with measurements in either fixed or live cells. Transmission is necessary for live cell work where the lasers must pass through coverglass and aqueous solution before interacting with a living cell. Live cells are grown on coverslips and mounted onto standard glass slides using a double-sided adhesive spacer with a pathlength of 5-10 µm to limit water background for copropagating (Fig. 2A-B). The chamber is filled with either PBS or media to ensure cell viability without IR interfering additives such as phenol red or protein and lipid found in fetal bovine serum. Counterprop mode necessitates a slightly more complex sample assembly than coprop mode. Because the lasers must be focused through objectives from both sides, a CaF_2_ coverslip must be mounted to another coverslip (glass or CaF_2_) rather than a glass slide, decreasing ease of handling and chamber construction (Fig. 2B). As long as a CaF_2_ coverslip is oriented towards the IR laser and Cassegrain objective (Fig. 2B), when a narrow spacer is used cells grown on either the top or bottom coverslip can be imaged. Growing on glass may be useful for cell adhesion. In our experience, imaging biological samples in counterprop with transmission collection has significantly lower signal to noise compared to the same spot collected in coprop transmission mode (Fig. 2C), which is consistent with our S/N analysis (Fig. S2). Therefore, our preferred live cell imaging modality is coprop transmission (Fig. 2A).

### Multispectral vs. hyperspectral imaging

Although hyperspectral images are data rich, due to the long collection times, on the order of hours, they are only compatible with fixed cells. Thus, live cell imaging necessitates multispectral imaging or the collection of few ad hoc full spectra. Hyperspectral data collected in fixed cells can be used to identify specific frequencies of interest for multispectral imaging and, for dynamic processes, to determine anticipated changes in IR absorbance between different conditions. Nevertheless, significant data is lost when information is only collected at 2-5 frequencies. Do multispectral images match hyperspectral images? How accurate is multispectral data processing at correcting for background and replicating trends seen with hyperspectral data?

To answer these questions, we compared hyperspectral and multispectral images of fixed differentiated 3T3-L1 cells fed ^13^C glucose. In differentiated 3T3-L1 cells, the ^13^C glucose is metabolized through glycolysis and then *de novo* lipogenesis over the span several days. The ^13^C glucose is primarily incorporated into triglycerides, which are stored inside of lipid droplets.^17,18^ The metabolic rate and spatial distribution can be tracked as a ratio of the ^13^C:^12^C ester carbonyl of triglycerides. Hyperspectral images were thresholded to only include pixels with significant native lipid signal; this was defined as an average intensity over the range of 1737-1757 cm^-1^ of at least 2.5% of the vector normalized spectrum. A ratio image was calculated at the remaining pixels as the average intensity of the labeled ^13^C=O lipid band (1693-1713 cm^-1^) over native ^12^C=O lipid band (1737-1757 cm^-1^) after correction for overlap with the amide-I band (Equation 1). A similar protocol was followed for the multispectral images except that single frequency images were collected at only the native and labeled lipid bands and the protein band, 1747, 1703, and 1655 cm^-1^, respectively (Equation 2) and a threshold of 5% of maximum lipid intensity was used. Multispectral images at the three frequencies could be collected in 5-15 minutes total for a pixel size of 250 nm while hyperspectral required 15-24 hours for a pixel size of 500 nm. We found qualitatively good agreement between the hyperspectral and multispectral ratio images (Fig. 3A-B).

**Figure 3.**
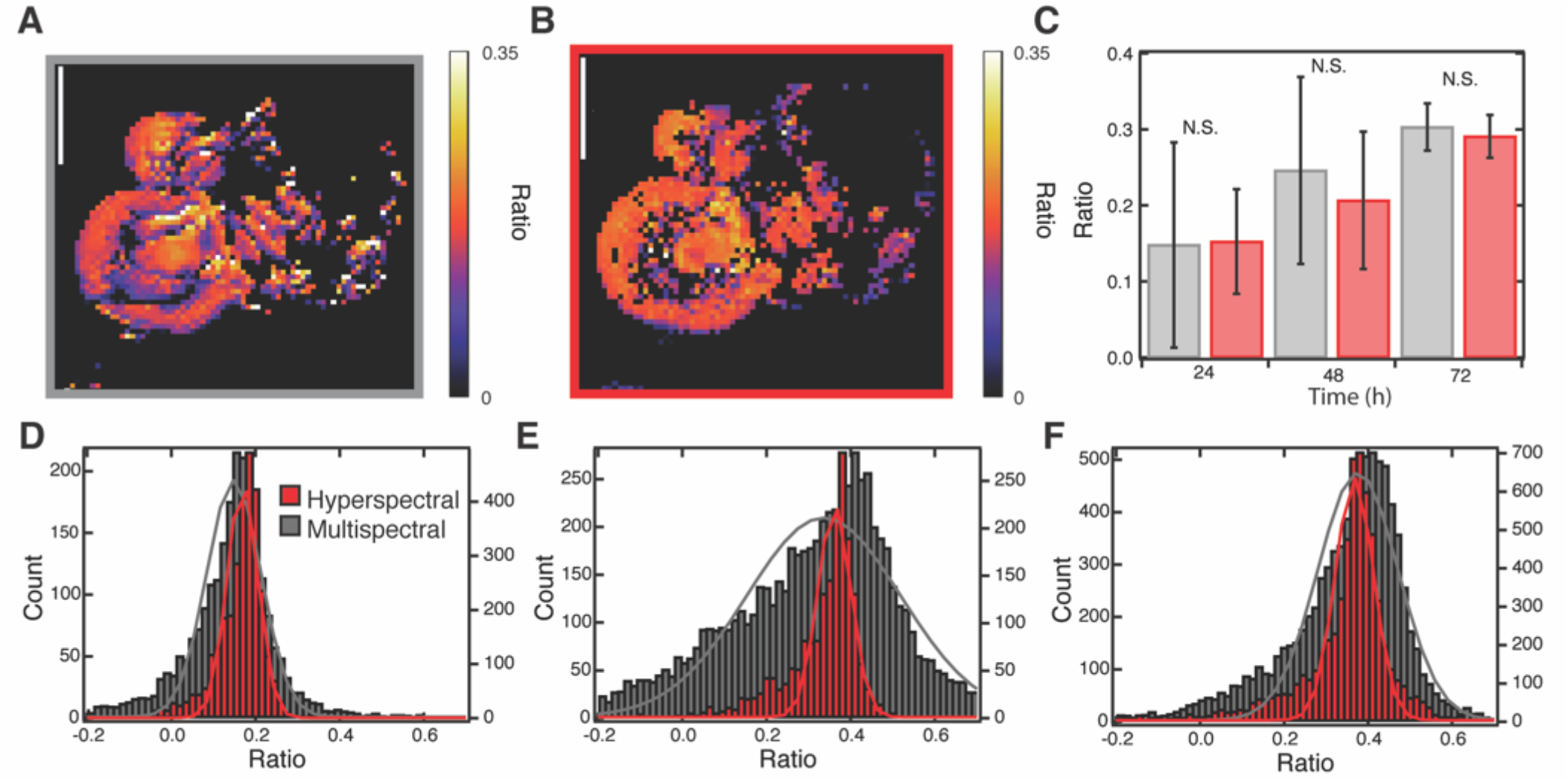
Hyperspectral and multispectral ratio imaging comparison. Representative (A) multispectral and (B) hyperspectral ratio image at 24 hours post ^13^C glucose feeding. Scale bars equal to 10 μm. A pixel size of 250 nm with a scan speed of 150 μm/s was used for the multispectral imaging and a step size of 500 nm was used for hyperspectral imaging. (C) Averages of the ratio of ^13^C lipid to ^12^C lipid for hyperspectral (red) and multispectral (grey) images (n=3 cells per timepoint). Histograms of the ratio of ^13^C lipid to ^12^C lipid and corresponding fits to a Gaussian distribution as calculated from hyperspectral (red, left axis) and multispectral (grey, right axis) images in three differentiated 3T3-L1 cells at (D) 24, (E) 48, and (F) 72 hours after feeding with ^13^C glucose. See methods for ratio calculation. Images collected using sequence method.

The average of three hyperspectral images of cells as compared to the three matching multispectral images showed good agreement (Fig. 3C) (p>0.15 at each timepoint in a paired t-test). Note, the high cell to cell variability in the small sample size (n=3), limited by the time required to collect hyperspectrals, somewhat obscures the metabolic trend of increased ^13^C over time that was observed with high confidence in larger multispectral datasets in our previous work.^17,18^ Nevertheless, the agreement between the hyperspectral and multispectral data in each cell and at each time point confirms the effectiveness of using multispectral imaging to rapidly collect larger sample sizes. However, a histogram of the ratios obtained at each pixel reveals systematic differences between the hyperspectral and multispectral images. Ratios generated from the multispectral images had a wider distribution than those calculated from the hyperspectral images (Fig. 3D-F). A possible explanation for the wider distribution of ratios in the multispectral data is that small peak shifts in the native lipid band (caused by differences in lipid composition across the cell) are detected in a single frequency multispectral image but not detected in the hyperspectral images which use the average frequency of a 20 cm^-1^ bandwidth. However, narrowing the average range for the hyperspectral image processing to 2 cm^-1^, similar to the ∼1.1 cm^-1^ laser bandwidth used for multispectral images, does not significantly increase the FWHM of the Gaussian (Fig. S3). Similarly, adjusting the spatial resolution via binning the multispectral image pixels to 500 nm to match the hyperspectral images does not influence width (Fig. S4).

Instead, differences between hyperspectral and multispectral imaging are caused by the increased scan speeds used in multispectral imaging and possibly error in the stage movement. Hyperspectral images collected at 500 nm resolution have a “scan speed” of less than 0.06 μm/s given that each spectrum takes approximately 8 s to collect. We suspect that this greatly reduces stage vibrations and other sources of error. To prove this, we conducted multispectral imaging of amide-II to amide-I ratio at a low scan rate on fixed Huh-7 cells and saw narrowing of the histograms (Fig. 4). However, imaging time was increased by a factor of 10. Together, this showcases that multispectral imaging is an excellent tool with speeds several orders of magnitude faster than hyperspectral imaging, albeit with slightly increased error. Multispectral imaging can therefore be reliably used for live cell imaging. Images can be collected by either the sequence or interleaving method. The interleaving method reduces any artifacts due to sample drift between images, but we observed no differences in ratio images of fixed cells (Fig. S5A-B). Pixel size, within the resolution of the instrument, also did not alter ratio images (Fig. S5B-D).

**Figure 4.**
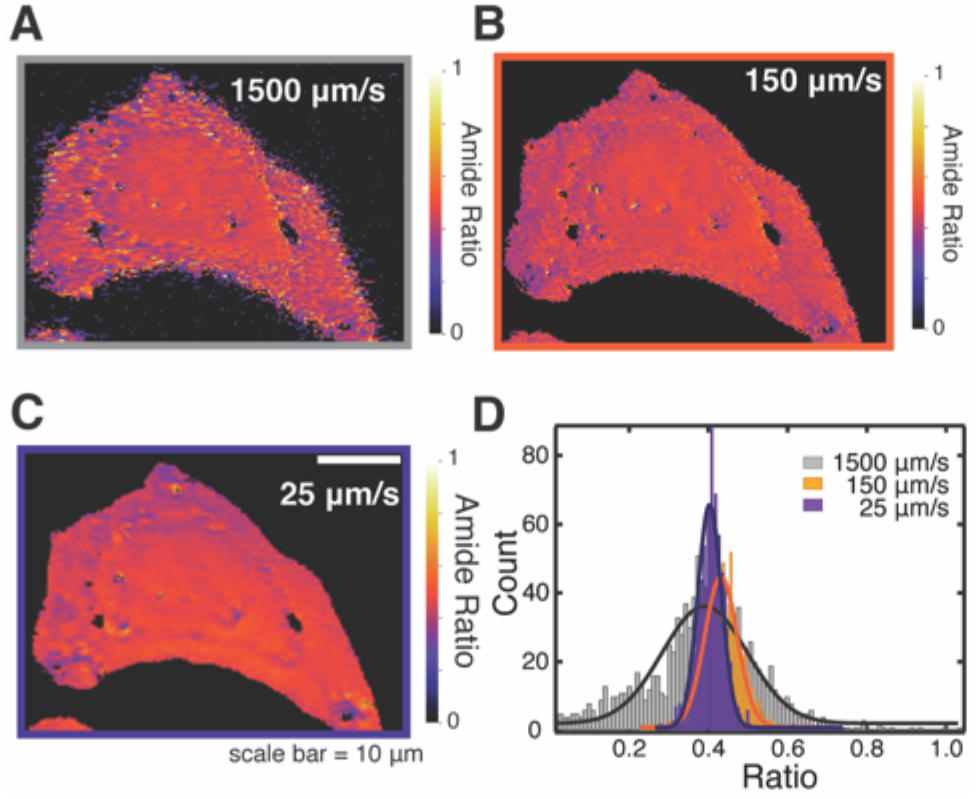
Scan speed determines noise. Fixed Huh-7 cell multispectral ratio images of amide-II (1540 cm^-1^) to amide-I (1655 cm^-1^) at scan speeds of (A) 1500 µm/s, (B) 150 µm/s and (C) 25 µm/s. Pixel size was 250 nm. (D) Histograms of the ratios of pixels shown in A-C and corresponding fits to a Gaussian distribution. Images collected using the sequence method.

### Validating infrared probes for OPTIR

While the intrinsic IR modes of biological molecules can be used to assign a variety of cellular components (Fig. S1), they lack the specificity to track specific molecules in the cellular milieu. Vibrational probes can be used to track a specific cellular process^3,4^ or biological molecule^37^ of interest and in some cases can even reveal local structural and/or environmental information. IR probes are most useful when there is not significant cellular background in the spectral region of the vibrational mode. This makes the cell silent window particularly attractive (Fig. 1A). Although IR probes have been developed and used for decades,^37,38^ proving their compatibility with a commercial OPTIR system in biological samples is not trivial. High spatial resolution means reduced probe concentrations in the focal volume, which could reduce visibility of the probe if concentrations are too low. Furthermore, because OPTIR uses both visible and IR lasers, probes must be photostable to both visible and infrared light.

Here we highlight the need to validate existing IR probes when using OPTIR using noncanonical amino acids with azide moieties (Fig. S6). The azide moiety has a strong infrared absorbance near 2100 cm^-1^ in the cellular silent region where there is no significant interference from other cellular biomolecules. Others have followed lipid^15^ and carbohydrate^19^ metabolism via azide labeled precursors with OPTIR. We demonstrate OPTIR detection of global and site-specifically labeled proteins using azide probes azidohomoalanine (AHA) and 4-azidomethyl-L-phenylalanine (AzmF), respectively, and importantly demonstrate the failure of OPTIR to detect site-specific incorporation of 4-azido-L-phenylalanine (AzF).

AHA is a methionine analog that is readily incorporated into proteins via traditional protein synthesis.^39^ By employing a methionine auxotroph in a methionine-free environment, all methionine residues in newly synthesized proteins should be replaced with AHA residues. After feeding, AHA signal in the cellular silent region was detectable in both *E. coli* and human U2OS cells (Fig. 5A, Fig. S7). Highlighting OPTIR sensitivity, we are able to visualize AHA in single fixed *E. coli* (Fig. S8). In the future, AHA can be used as a general probe of protein synthesis and degradation.

**Figure 5.**
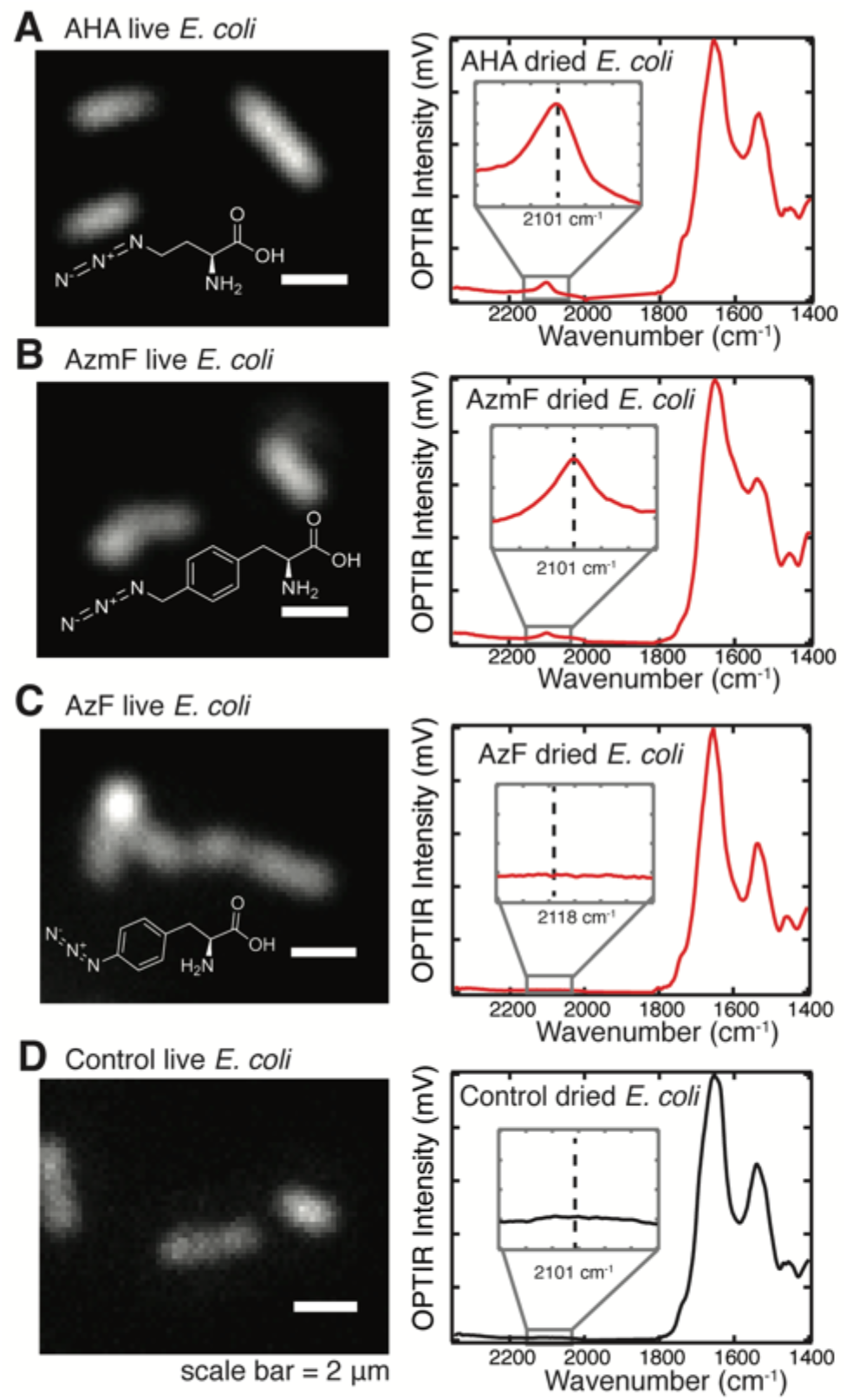
Azide probes in *E. coli*. Left: Fluorescent images of live *E. coli* in PBS overexpressing (A) AHA labeled sfGFP, (B) sfGFP 150AzmF, (C) sfGFP 150AzF, and (D) unlabeled sfGFP. Insets show probe structure. Right: Representative OPTIR spectra of dried, bulk *E. coli* overexpressing (A) AHA labeled sfGFP, (B) sfGFP 150AzmF, (C) sfGFP 150AzF, and (D) unlabeled sfGFP. Insets highlight cell silent region where azide moieties absorb.

Alternatively, the azide probe AzmF can be incorporated site-specifically into a protein of interest via amber codon suppression.^40^ We incorporated AzmF into overexpressed sfGFP at position 150 in *E. coli* using amber codon suppression.^41^ These cells were fluorescent, confirming successful AzmF incorporation as otherwise the protein would be truncated and unable to fold and fluoresce. Despite only being incorporated into a single location of a single, overexpressed protein, the AzmF signal was easily detectable in the cell silent region (Fig. 5B). In the future, this handle can be used to map the location of the labeled protein, and as the IR signal of AzmF is sensitive to local environment, this probe may serve as a tool for protein localization and structural determination.^25^

However, incorporation of an azide probe does not always mean detection. AzF, another unnatural amino acid that can be incorporated using amber codon suppression, is incorporated into *E. coli* expressing sfGFP 150AzF, as evidenced by fluorescence (Fig. 5C). Yet the OPTIR signal in the cell silent region is not different from control *E. coli* (Fig. 5D). This is notable since AzF has been used as an IR probe before.^42^ We posit that this is due to the visible laser used in OPTIR causing loss of the azide bond as AzF is photoactivatable while AzmF is not.^25^

### Live cell imaging of OPTIR probes

Spectra collected in live cells share many features with those collected in fixed cells with the addition of a dominant water band (Fig. S9). The dominant water band can be corrected for in multispectral ratio images via a straightforward linear correction, although complete preservation of all structural information contained in the amide-I band is not possible.^18^ We have previously found good agreement between ^13^C/^12^C lipid ratios in fixed and live Huh-7 and 3T3-L1 cells.^17,18^ Alternatively, to avoid overlap with biomolecules in the cellular background IR probes that make use of the cell silent region can be employed. However, it is worth noting that the relatively weak bend + libation water combination band (centered at 2130 cm^-1^) can complicate live cell measurements of IR probes in the cell silent region. Therefore, not all IR probes that are validated in fixed cells for OPTIR can be easily translated to live cells.

It is also important to note that the same probe may be primarily used for different metabolic processes in different cell lines, enabling (and limiting) the tracking of different metabolic processes in different cell lines and organisms. For example, we have previously used isotopic precursor ^13^C glucose to track lipogenesis via incorporation of the isotope label into cellular lipids.^18,43^ Feeding ^13^C glucose leads to shifts across the cell’s infrared profile where ^13^C is incorporated. The heavier isotope causes a redshift in vibrations when present as frequency is inversely proportional to reduced mass according to Equation 5:^44^

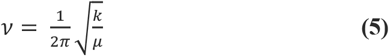

where ν is frequency, k is the force constant of the bond, and μ is reduced mass. Shifts in the spectrum, i.e. what biomolecules ^13^C is incorporated into from ^13^C glucose, depend on the metabolism of the cell line in question.

When ^13^C glucose is fed to *E. coli* cells, the bulk of the carbon is used for protein synthesis. Highlighting this, we show that *E coli* grown in ^13^C glucose minimal media have a shifted amide-I and II band and, furthermore, that the amide-II band can be used to visualize live single *E. coli* in H_2_O based buffer (Fig. 6A, Fig. S10A). ^13^C glucose metabolism can be used to track susceptibility of *E. coli* to antibiotics and application of live cell OPTIR measurements would remove time consuming fixative processes.^45,46^

**Figure 6.**
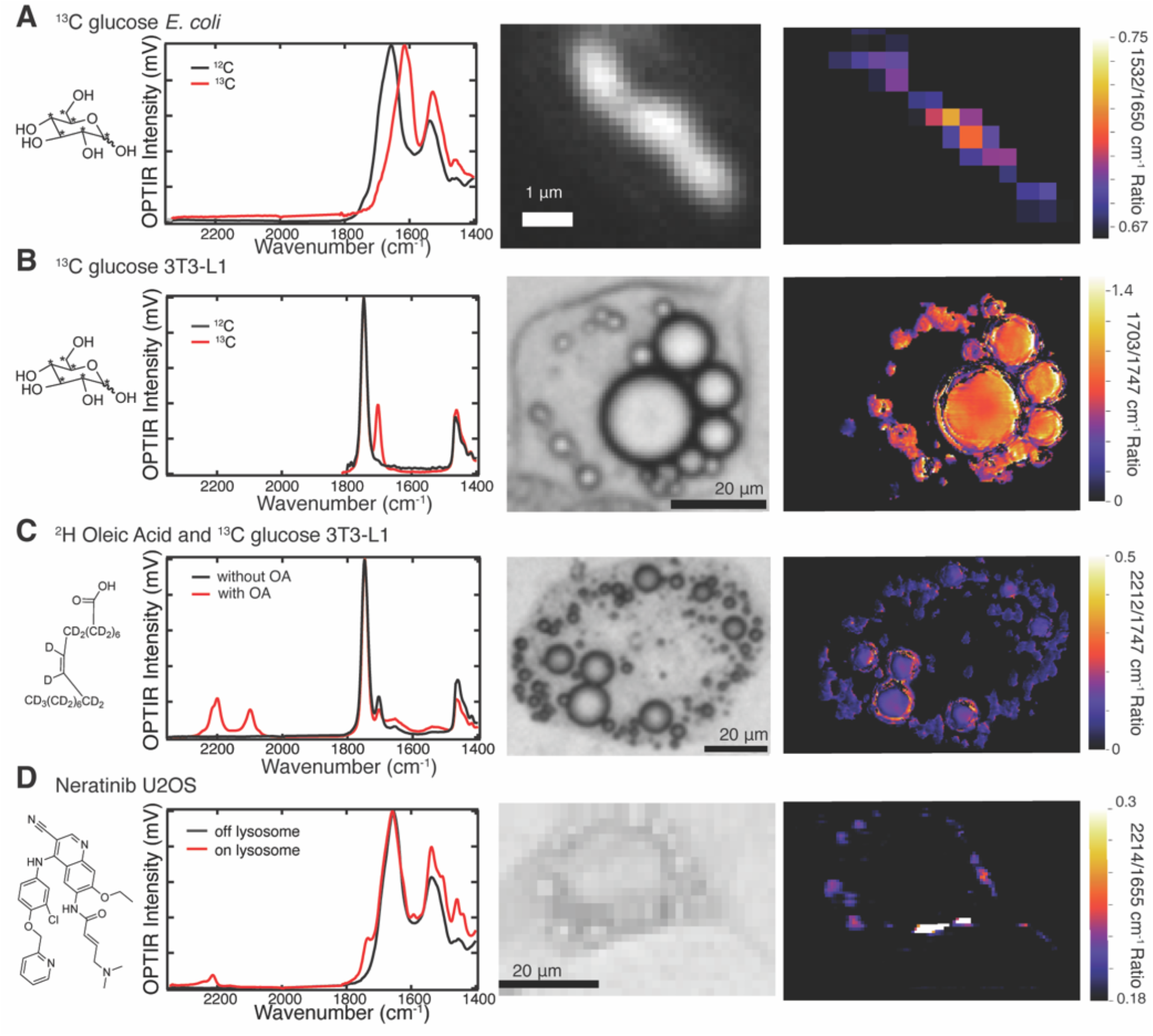
Metabolic probes in biological systems. Left: (A) Structure of ^13^C glucose where asterisks represent ^13^C and average spectra of fixed *E. coli* after feeding ^12^C glucose media (black) or ^13^C glucose media (red). (B) Structure of ^13^C glucose where asterisks represent ^13^C representative spectra of a fixed differentiated 3T3-L1 cell with (red) and without (black) feeding of ^13^C glucose (72 hrs, 4.5 g/L). The control spectrum was collected on a previous mIRage OPTIR model.^17^ No data in cell silent region. (C) Structure of deuterated oleic acid and representative spectra in a fixed differentiated 3T3-L1 cell fed ^13^C glucose (24 hrs, 4.5 g/L) with (red) and without (black) feeding deuterated oleic acid (24 hrs, 60 μM). (D) Structure of neratinib and representative spectra of a fixed U2-OS cell fed neratinib (7 hrs, 5 μM) on (red) and off (black) a lysosome. Center: (A) Fluorescent image of single live *E. coli* expressing sfGFP after feeding ^13^C glucose media, (B) bright field image of live differentiated 3T3-L1 cell with feeding of ^13^C glucose (72 hrs, 4.5 g/L), (C) bright field image of live differentiated 3T3L1 cell with feeding deuterated oleic acid (24 hrs, 60 μM) and ^13^C glucose (24 hrs, 4.5 g/L), and (D) bright field image of live U2-OS cell fed neratinib (7 hrs, 5 μM). Right: OPTIR ratio images of (A) 1532/1650 cm^-1^ ratio of single live *E. coli* expressing sfGFP after feeding ^13^C glucose media, no threshold used, (B) corrected 1703/1747 cm^-1^ ratio of live differentiated 3T3-L1 cell with feeding of ^13^C glucose (72 hrs, 4.5 g/L) (corrected with equation 4), (C) corrected 2212/1747 cm^-1^ ratio of live differentiated 3T3-L1 cell with feeding deuterated oleic acid (24 hrs, 60 μM) and ^13^C glucose (24 hrs, 4.5 g/L) (corrected with equation 3) (D) 2214/1655 cm^-1^ ratio of live U2-OS cell fed neratinib (7 hrs, 5 μM) no threshold used. All ratio images collected using the sequence method except for ^13^C E coli, which were collected using the interleaved method.

Conversely, in mammalian fat (differentiated 3T3-L1) and liver (Huh-7)^18^ cells the bulk of the glucose is directed to *de novo* lipogenesis, leading to the appearance of peaks associated with ^13^C labeled lipids (Fig. 6B, Fig. S10B). The most notable of these is the red-shifted lipid ester carbonyl stretch at 1703 cm^-1^. Interestingly, the unlabeled lipid ester carbonyl stretch appears at slightly different positions in Huh-7 (1744 cm^-1^) and differentiated 3T3-L1 (1747 cm^-1^) cells hinting at different lipid compositions (Fig. S1). Upon labeling, the lipid ester carbonyl peak shifts to 1703 cm^-1^ for both cell types suggesting that *de novo* lipogenesis over the time frame studied (72 hours) leads to more homogenous lipids, likely triacylglycerol (Fig. 6B, Fig. S10B).^47^ Live imaging is necessary to preserve lipid morphology and reveals rates of DNL with high spatial resolution across the cell.^43^

In the spirit of multicolor fluorescent imaging, IR probes can be multiplexed together to track multiple processes in parallel.^48^ In Fig. 6C we couple ^13^C glucose feeding with ^2^H oleic acid feeding in 3T3-L1 cells. C-D stretches appear in the cell silent region and deuterated oleic acid uptake and esterification can be visualized in living adipocytes over the background of cells fed ^13^C glucose alone (Fig. 6C, Fig. S10B). We^18^ and others,^49^ have used deuterated fatty acids to track metabolic pathways such as fatty acid scavenging and esterification with OPTIR.

Finally, certain molecules contain IR active moieties that appear in the cellular silent region without the need for probe incorporation. We demonstrate that neratinib, a cancer drug known to be trapped in the lysosome, can be visualized with OPTIR using the intrinsic nitrile stretch in live U2OS cell lysosomes (Fig. 6D, Fig. S10C). This has been previously observed by Raman microscopy of fixed cells.^50^ Many drugs contain nitrile or azide moieties and OPTIR may be a useful tool for live cell, spatially resolved pharmacokinetics.

### Organism imaging and future directions

OPTIR is also compatible with whole organism imaging.^12^ We predict that dynamic measurements taken via multispectral imaging and the flexibility and non-perturbative nature of IR probes will allow for better understanding of in organism metabolism and catabolism. Here we demonstrate that OPTIR can be collected in two model organisms, *H. exemplaris* tardigrades and zebrafish embryos (Fig. 7, Fig S11). Ratio imaging in tardigrade reveals consistent amide-II to water signal indicating that proteins are evenly distributed throughout the organism (Fig. 7C, left); Conversely, ratio images of lipid to water highlight areas of increased lipid, possibly corresponding to algal growth on and next to the tardigrade (Fig. 7C, right). Future studies could examine protein synthesis during tardigrade desiccation or changing IR signals in the embryo as its cells differentiate. IR probes could be used to follow metabolism in these organisms or localize specific proteins without the disruption caused by fluorescent tags.^3^

**Figure 7.**
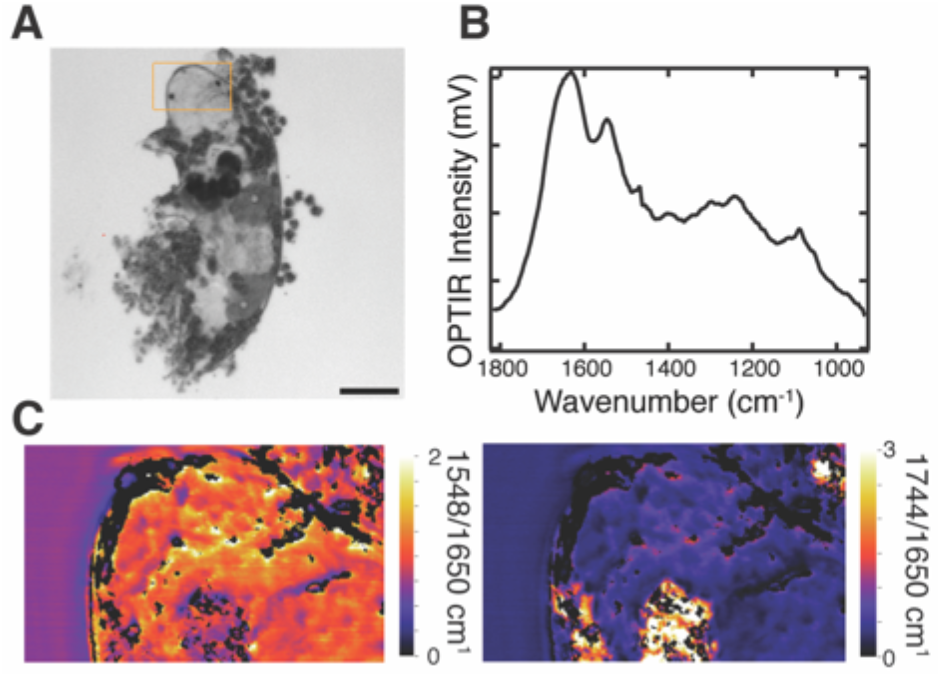
Live organism OPTIR. (A) Live tardigrade brightfield image. Scale bar is 50 μm. (B) Single OPTIR spectrum collected in live tardigrade and (C) OPTIR ratio images of 1548/1650 cm1 (left) and 1744/1650 cm-1 (right). Imaged area outlined in orange in (A). Collected in coprop transmission. Images collected using sequence method.

## CONCLUSION

This work demonstrates the flexibility of OPTIR as a tool for understanding biological systems. We have been able to showcase a variety of collection methods on a commercial instrument and confirm that multifrequency imaging collected in sequence at carefully selected frequencies can provide information equivalent to hyperspectral imaging at 20-30x the speed (minutes compared to 10-20 hours). This allows for detailed live cell imaging. Moreover, the application of IR active probes allows for investigation of specific proteins and metabolic pathways. Together, these advancements make OPTIR a powerful technique for cellular systems and introduces the possibility for future work to bring these probes and this dynamic imaging into whole organisms.

## ASSOCIATED CONTENT

## Supporting Information

The Supporting Information is available free of charge on the ACS Publications website.

Supplemental methods of AHA incorporation and zebrafish imaging, supplemental figures of average cell OPTIR spectra, controls for hyperspectral and multispectral imaging, azide probe OPTIR spectra and images, live cell and live cell probes OPTIR spectra, and zebrafish OPTIR spectra. (PDF)

## AUTHOR INFORMATION

### Corresponding Author

* Caitlin M. Davis

### Notes

The authors declare no competing financial interest.

## ACKNOWLEDGMENTS

This work was supported by the Arnold and Mabel Beckman Foundation. Hepatocyte research was conducted while Caitlin Davis was a Hevolution/AFAR New Investigator Awardee in Aging Biology and Geroscience Research. Adipocyte and zebrafish research was supported by National Institutes of Health (NIH) grant R35 GM151146. Tardigrade research was supported by NSF CAREER award MCB-2338323. S. O. S. and A. E. C. were partially supported by the NIH under training grant T32 GM008283. S. O. S. was partially supported by a National Science Foundation Graduate Research Fellowship under grant DGE-2139841. The authors thank Brahmmi Patel and Cailin Hoang for providing the zebrafish embryos and Marisa Barilla for tardigrade culturing.

---

Insert Table of Contents artwork here

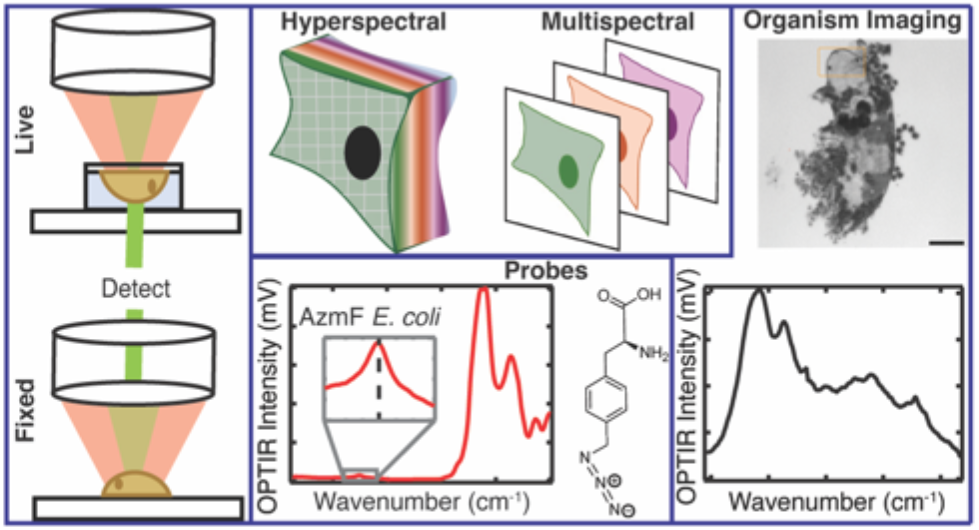

