## Supplementary Information for "Optical photothermal infrared imaging using metabolic probes in biological systems"

**Table of Content**

**Methods**…………………………………………………………………………………………………….. S2

**References**….……………………………………………………………………………………………… S2

**Figure S1.** Full assignments of different cell lines. ………………………………………….... S3

**Figure S2.** Root mean square noise of OPTIR modes of collection……………………….. S4

**Figure S3.** Limiting hyperspectral images to 2 cm^-1^ does not make them match multispectral images. …………………………………………………………………………………………………….… S5

**Figure S4.** Adjusting the spatial resolution via binning the multispectral image pixels does not have an effect on multispectral ratio histograms………………………………………….. S6

**Figure S5.** Sequence vs. interleaving image collection and pixel size within the resolution of the instrument does not affect multispectral ratios……………………………………………. S7

**Figure S6.** OPTIR spectra of azide probes…………………………………………………………. S8

**Figure S7.** AHA in fixed U2OS cell. …………………………………………………………………. S9

**Figure S8.** Single fixed *E. coli* AHA imaging..…………………………………………………..…. S10

**Figure S9.** Fixed vs live Huh-7 averages..……………………………………..………………..…. S11

**Figure S10.** Live cell spectra of probes....……………………………………..………………..… S12

**Figure S11.** Zebrafish OPTIR. ....……………………………………………..…..………………..… S13

**Methods**

Azidohomoalanine (AHA) incorporation in mammalian cells. U2OS cells were trypsinized and plated onto CaF_2_ coverslips. After 6 hours, media was exchanged for methionine, glutamine, and cystine free high-glucose DMEM (Gibco) supplemented with glutamine (4 mM), cystine (0.2 mM) and AHA (5 mM). After 24 hours, the cells were fixed and imaged.

Zebrafish imaging. Zebrafish embryos were selected 48 hours after fertilization and anesthetized using 0.4% tricaine solution (Ward’s Science, Rochester, NY). Embryos were placed directly on a microscope slide in 20 μL of E3 medium with tricaine^1^ and imaged in epi mode.

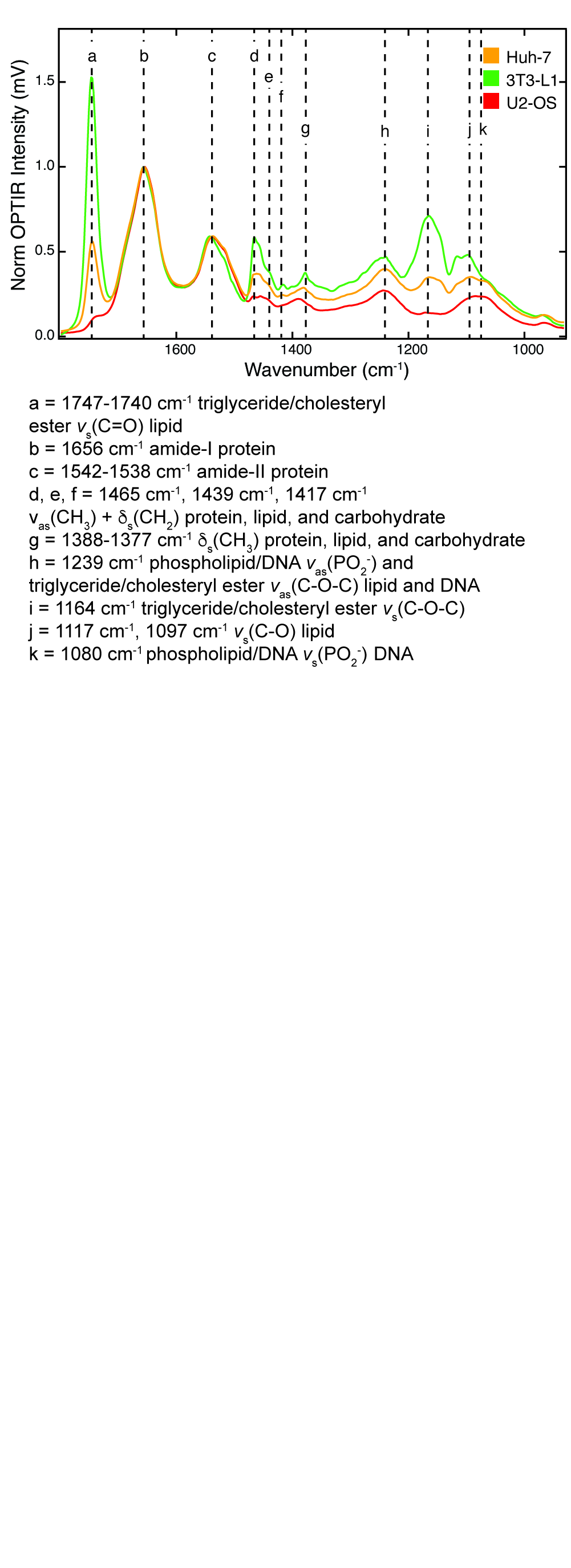

**Figure S1.** Full assignments of different cell lines. Average fixed cell spectra of Huh-7 (yellow), differentiated 3T3-L1 (green), and U2OS (red) generated by averaging a full hyperspectral image of one cell per cell line. Hyperspectral spacing of 500 nm. 3T3-L1 data collected on a previous microscope model, a mIRage IR microscope (Photothermal Spectroscopy Corporation, Santa Barbara, CA) integrated with a three-module-pulsed quantum cascade laser (QCL) system (Daylight Solutions, San Diego, CA) with a tunable range from 803 cm^-1^ to 1799 cm^-1^, as previously described.^2^

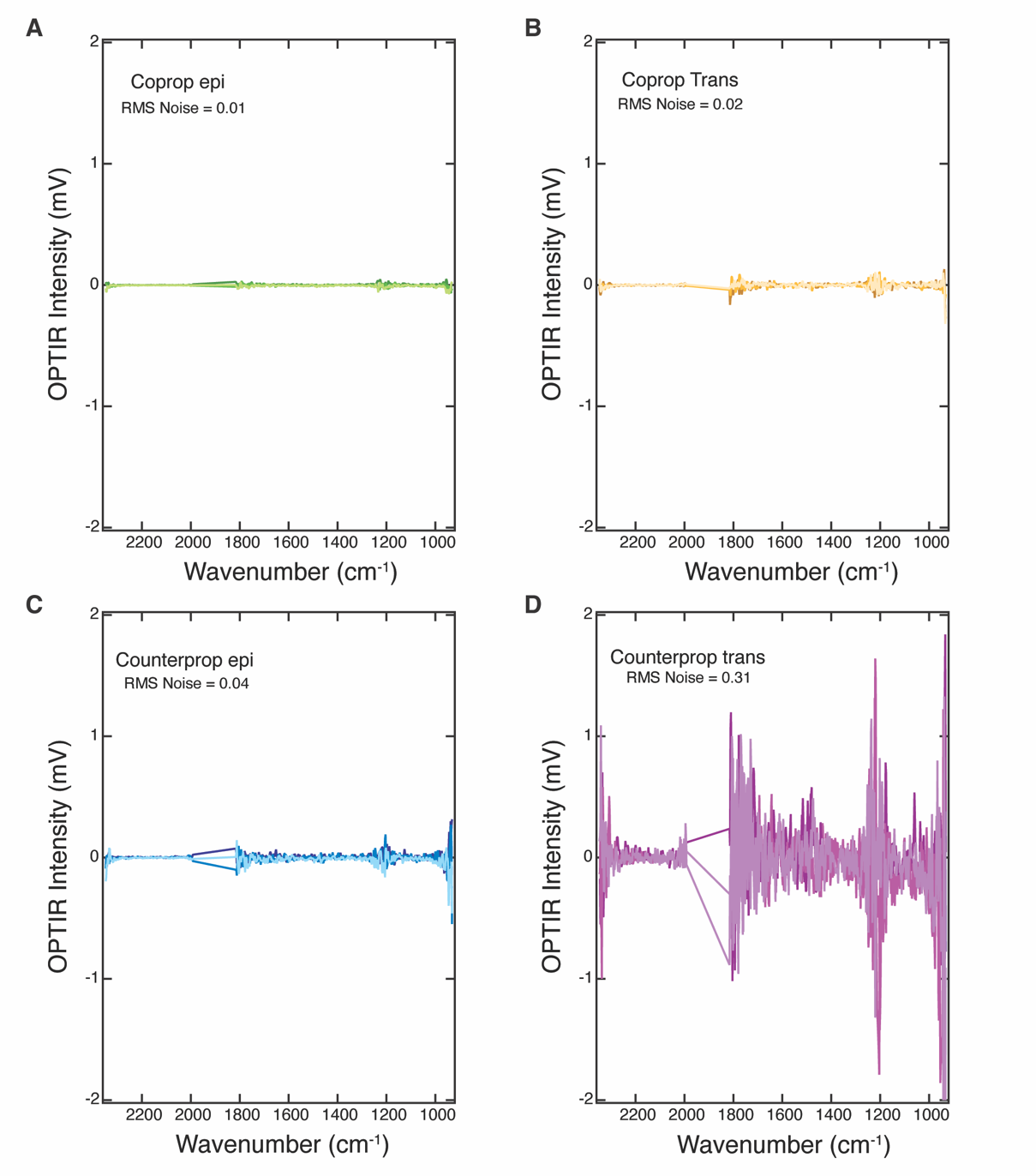

**Figure S2.** Root mean square noise of OPTIR modes of collection. Three acquisitions of an empty CaF_2_ coverslip taken in (A) copropagation epi (B) copropagation transmission (C) counterpropagation epi and (D) counterpropagation transmission. Note there is no coverage of 1816-1992 cm^-1^. Three acquisitions, IR power of 20%, probe power of 11% and detector gain of 1x was used.

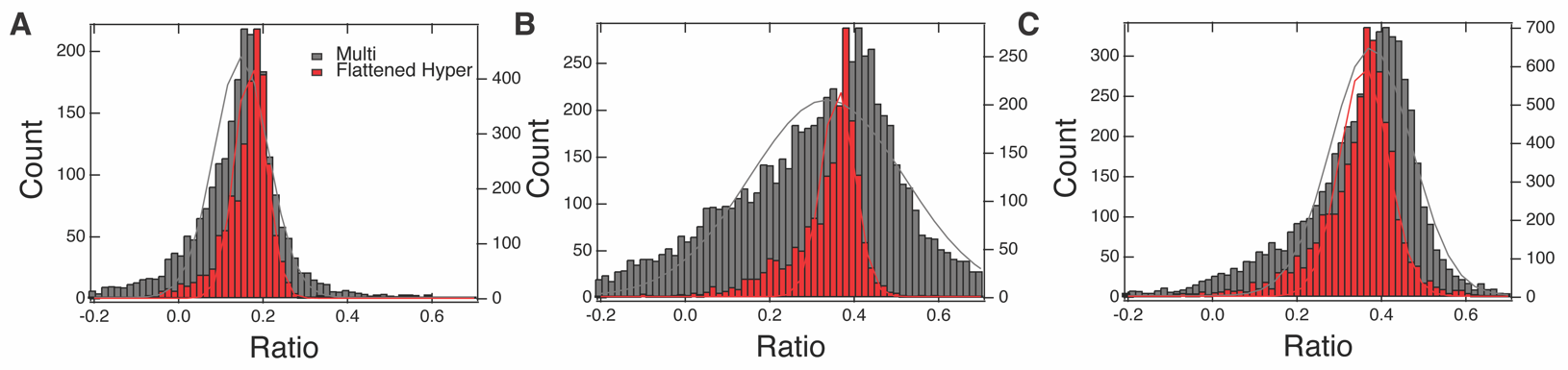

**Figure S3.** Limiting hyperspectral images to 2 cm^-1^ does not make them match multispectral images. Histograms of the ratio of ^13^C lipid to ^12^C lipid as calculated from hyperspectral images where only 2 cm^-1^ were used for ratio calculation (red, left axis) and multispectral (grey, right axis) images in three 3T3-L1 cells at (A) 24, (B) 48, and (C) 72 hours after feeding with ^13^C glucose. See methods for ratio calculation. Histograms overlaid with a fit to a Gaussian distribution.

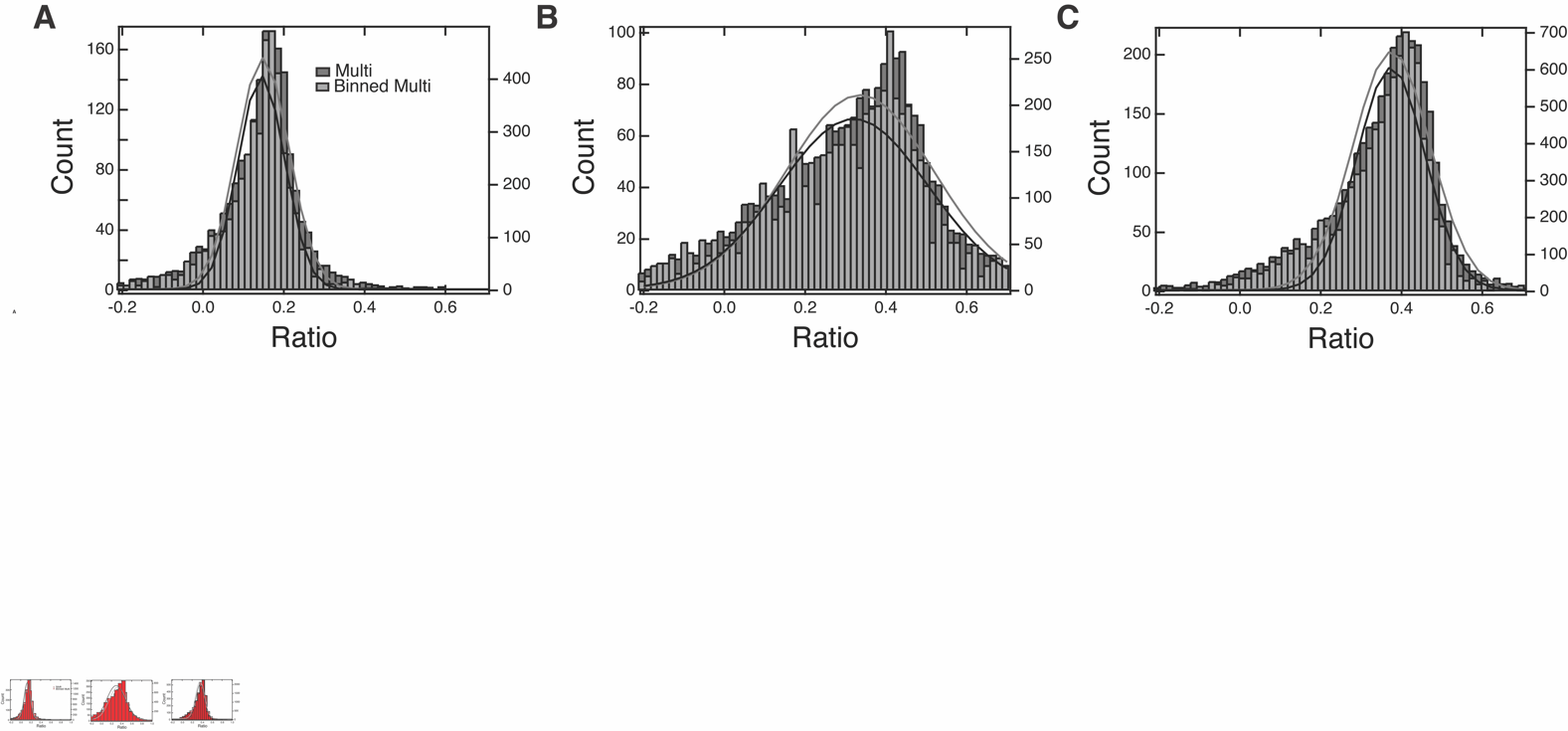
**Figure S4.** Adjusting the spatial resolution via binning the multispectral image pixels does not have an effect on multispectral ratio histograms. Histograms of the ratio of ^13^C lipid to ^12^C lipid as calculated from multispectral (dark gray, right axis) images and 2x2 binned multispectral (light gray, left axis) images in three 3T3-L1 cells at (A) 24, (B) 48, and (C) 72 hours after feeding with 13C glucose. See methods for ratio calculation. Histograms overlaid with a fit to a Gaussian distribution.

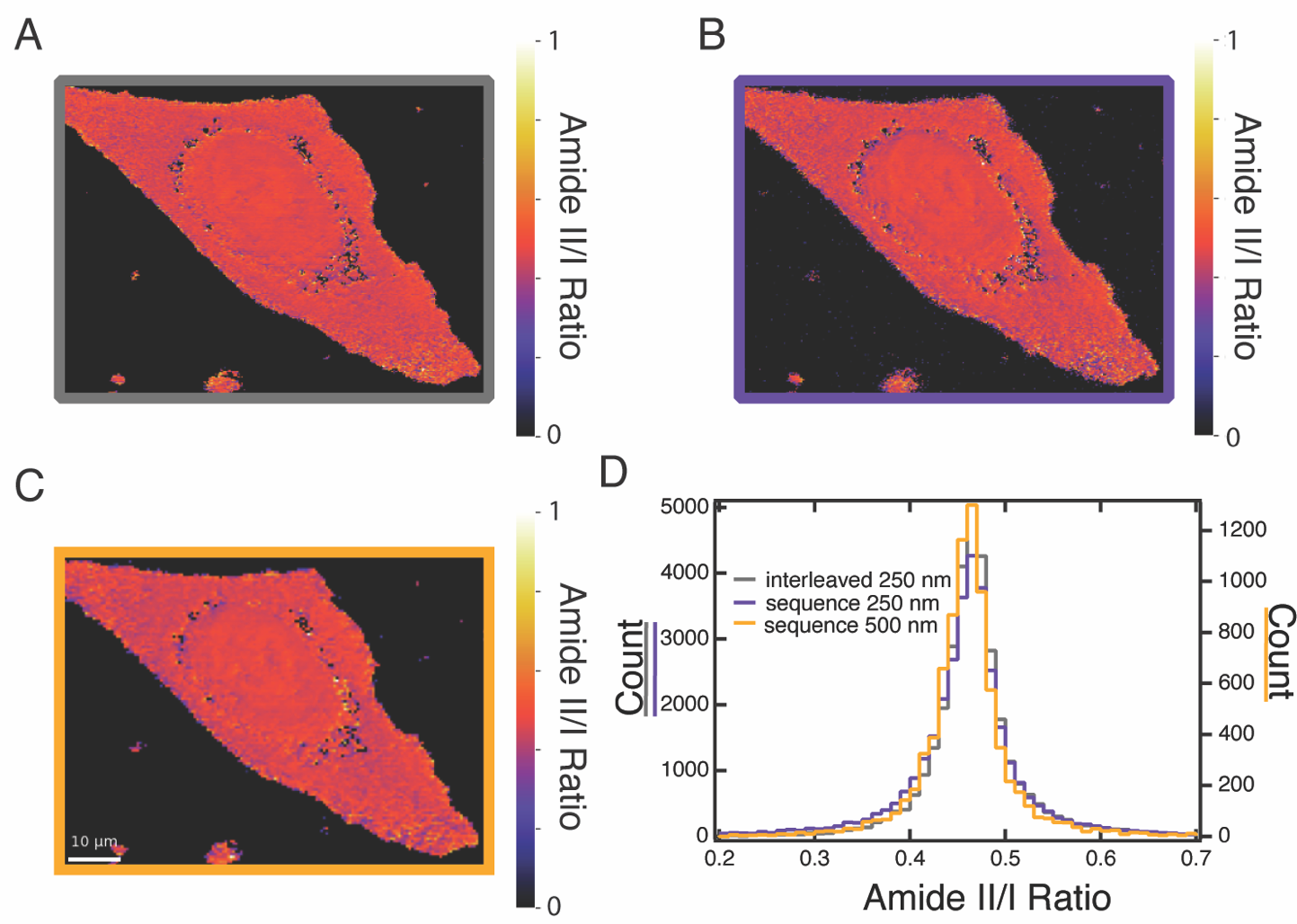

**Figure S5.** Sequence vs. interleaving image collection and pixel size within the resolution of the instrument does not affect multispectral ratios. Fixed U2-OS cell multispectral ratio images of amide-II (1540 cm^-1^) to amide-I (1655 cm^-1^) collected (A) using the interleaved method at 250 nm pixel size and the sequence method at (B) 250 nm and (C) 500 nm pixel size. (D) Histograms of the ratios of pixels shown in A-C. Left axis 250 nm pixel size and right axis 500 nm pixel size.

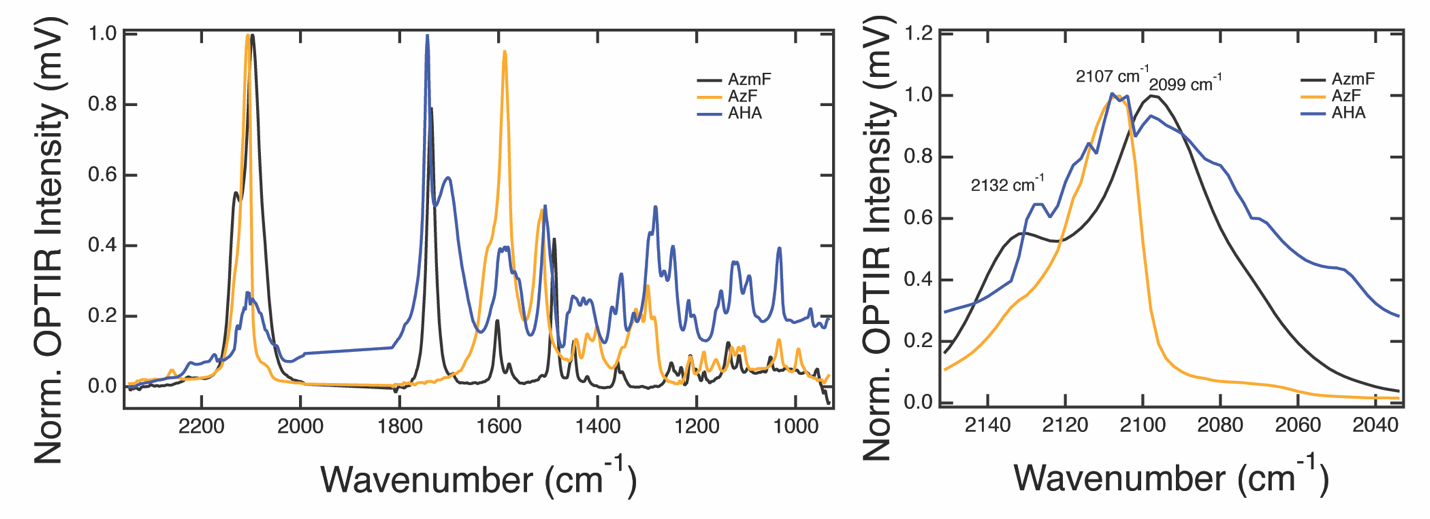
**Figure S6.** OPTIR spectra of azide probes. (left) Normalized OPTIR spectra of AzmF (black), AzF (orange), and AHA (blue). (right) Normalized azide region of the same spectra: AzmF (black), AzF (orange), and AHA (blue). Spectra collected in epi coprop from hydrated films of AzmF and AHA. Spectra of AzF is collected as a powder, as it is minimally soluble and photoreactive.

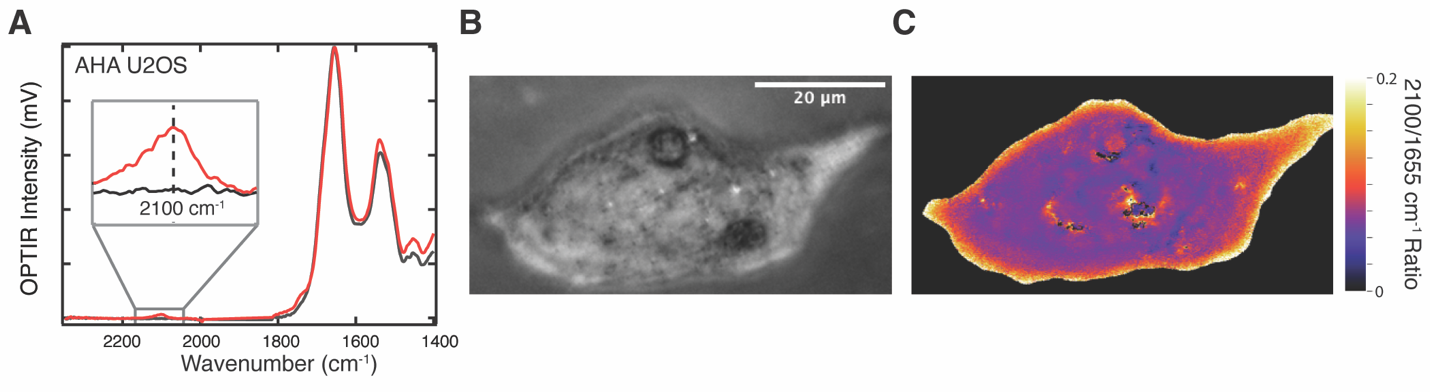

**Figure S7**. AHA in fixed U2OS cell. (A) Representative spectrum in a fixed U2OS cell with (red) and without (black) feeding AHA (24 hrs, 5 mM). Inset: Azide region. (B) Brightfield and (C) OPTIR ratio image (2100/1655 cm^-1^) of fixed U2OS cell incubated with AHA (24 hrs, 5 mM). Threshold of 2% at 1655 cm^-1^ used with no correction. Images collected using sequence method.

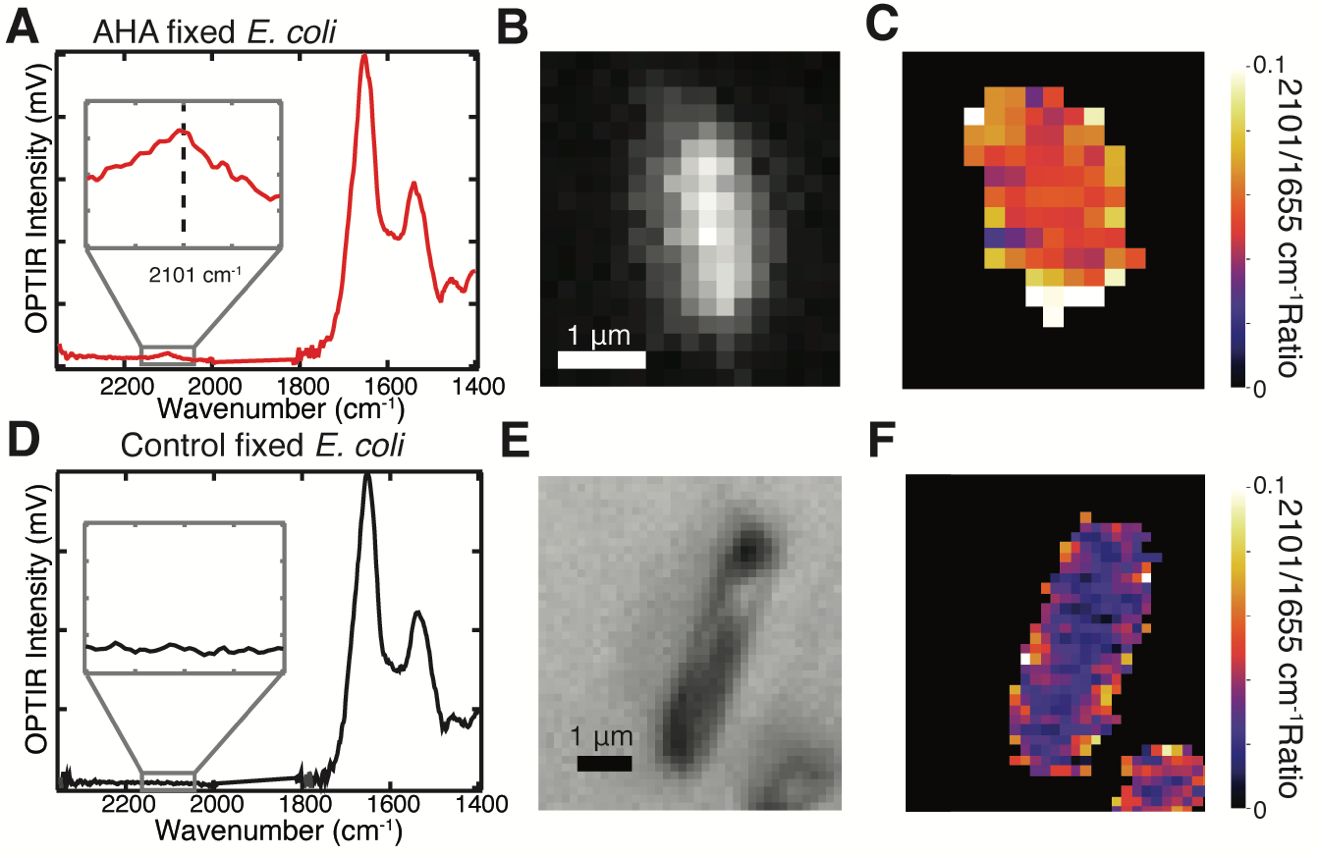

**Figure S8.** Single fixed *E. coli* AHA imaging. (A) OPTIR spectrum (8 accumulations), (B) fluorescent image, and (C) ratio image of single *E. coli* cell expressing AHA labeled sfGFP. (C) OPTIR spectrum (8 accumulations), (C) brightfield image (50X objective), and ratio image of single control *E. coli.* Threshold of 10% of 1655 cm^-1^ signal used for ratio image.

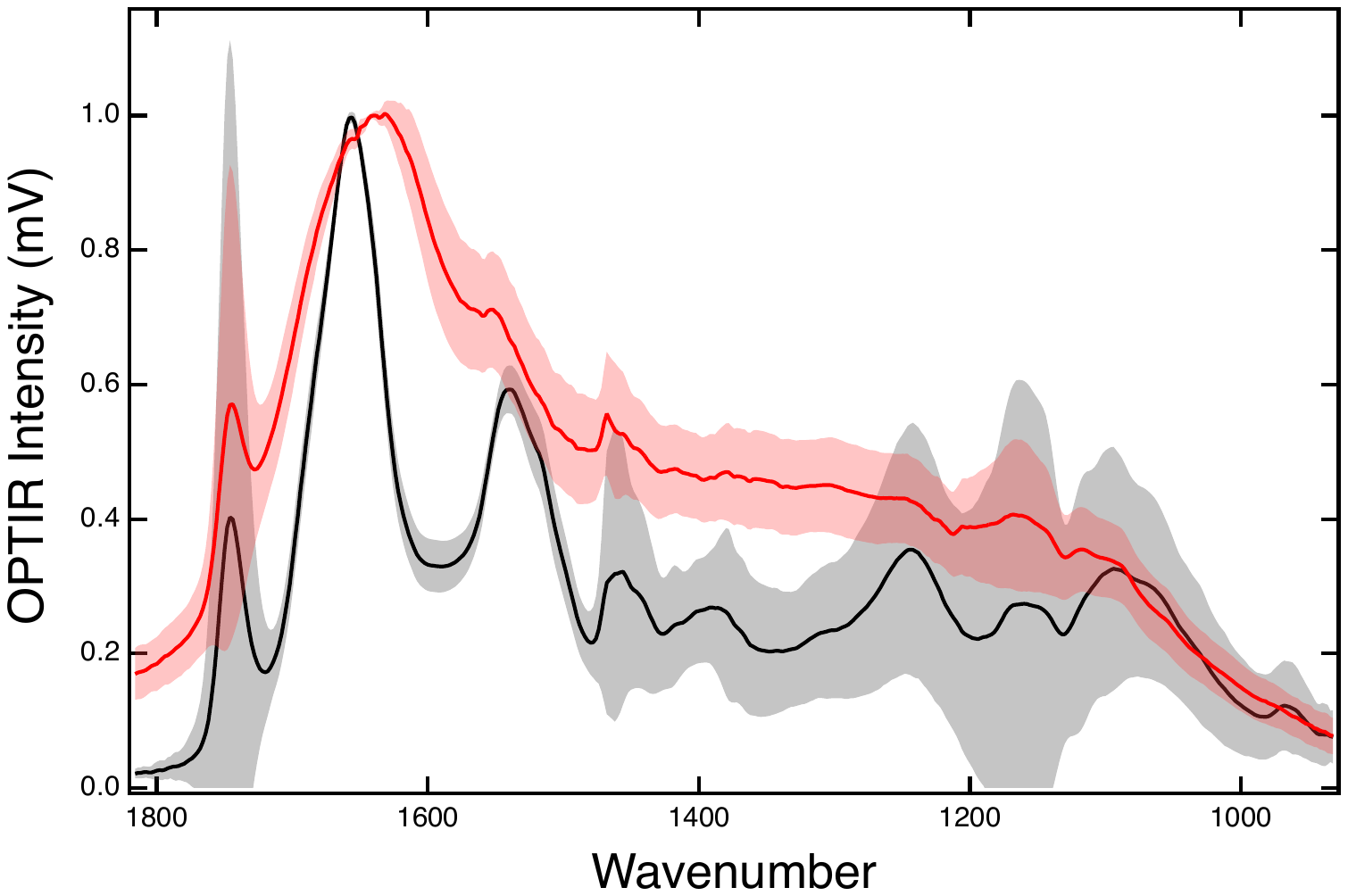

**Figure S9.** Fixed vs live Huh-7 averages. Averages of live (red) and fixed (black) Huh-7 cell spectra (n=16 cells).

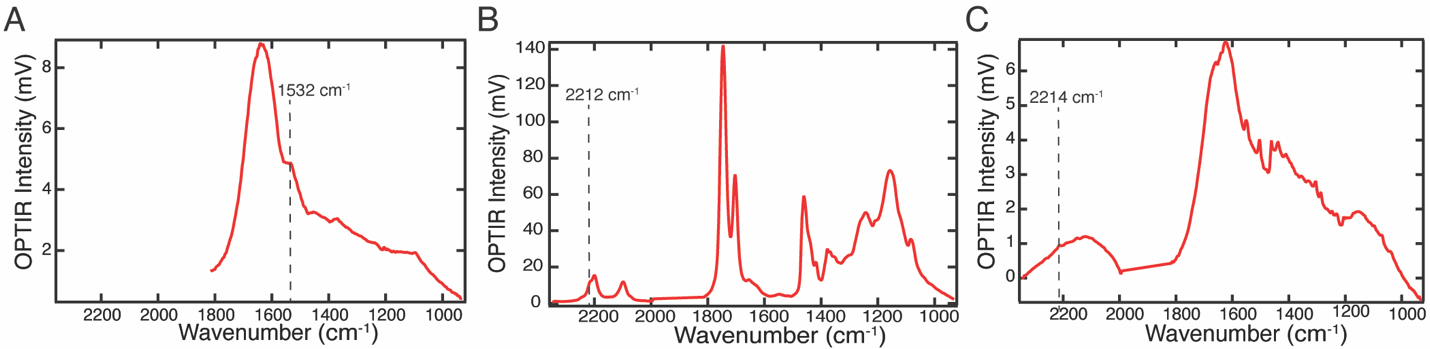
**Figure S10.** Live cell spectra of probes. Representative OPTIR spectra of (A) live *E. coli* expressing sfGFP after feeding ^13^C glucose media, (B) live differentiated 3T3-L1 cell fed deuterated oleic acid (60 μM) and ^13^C glucose (4.5 g/L) for 24 hours, and (C) live U2OS cell fed neratinib (7 hrs, 5 μM). Spectra collected in coprop transmission.

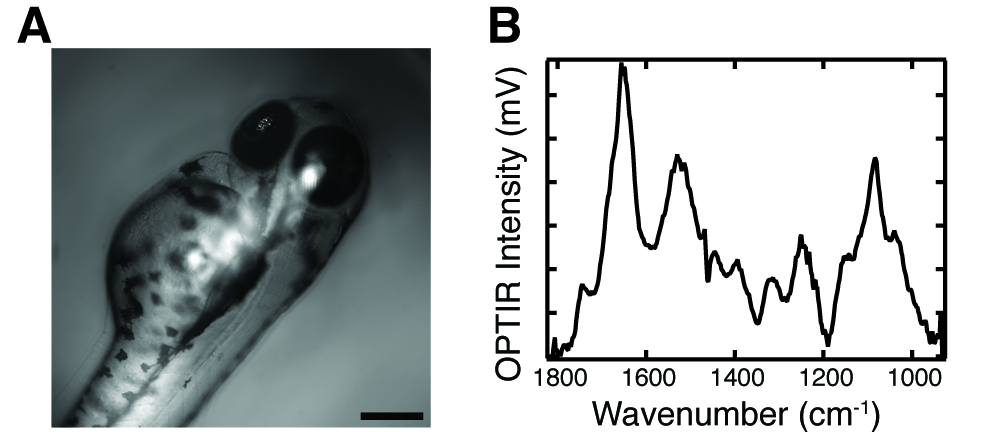

**Figure S11.** Zebrafish OPTIR. (A) Brightfield OPTIR microscope image of zebrafish embryo 48 hours post fertilization. Scale bar is 50 μm. The large size of the zebrafish embryo (~500 µm) precluded transmission OPTIR measurements, which are limited to short pathlengths due to water absorption in the mid IR. (B) Single OPTIR spectrum of the zebrafish embryo in a droplet of water collected in coprop epi mode. This was possible as a larger volume of water is required to encapsulate zebrafish embryos than cultured cells, slowing the drying process.
